# Identification and in silico comparison of transposable elements of Solanum tuberosum subsp. andigena from two localities of Peru

**DOI:** 10.1101/2021.08.03.453701

**Authors:** Sheyla Carmen Sifuentes, Victor Cornejo Villanueva

## Abstract

The potato is one of the main crops in Peru, being one of the pillars in the country’s economy. It also has a cultural importance, being a basic element of Peruvian gastronomy and an emblematic crop of the Peruvian highlands, where small farmers over time have been able to conserve and cultivate 6048 native varieties. In Peru, around 300,000 hectares of potatoes are planted annually, and in 2019 Peru remained the main potato producer in Latin America, registering an annual production of 5.3 million tons. Therefore, it is necessary to study and characterize the genome of this very important tuber for the country. Although there are many potato genomic studies in the world and in Peru; these have been based on the search for genes of importance, genetic diversity, etc. and little has been deeply studied about the transposable elements (ETs) that are part of its genome. It is known that in many eukaryotes, transposable elements are powerful drivers of genome evolution, and they occupy a considerable part in plant genomes, so with these studies, more could be understood about how these ETs have intervened in the evolution of the potato in Peru; thus, how these mechanisms would help in the future to the survival of the species in scenarios of climate change, among others. This study proposes the identification and comparison of the transposable elements of the only 2 complete genomes of potatoes grown in Peru, found in the NCBI database, one from the department of Puno and the other from Cusco. There are various approaches to search for ETs in genomes, but for a complete and reliable annotation, it has been shown that the best strategy is the adoption of combined approaches, in this study, we will use 2 approaches, one based on homology, with the program RepeatMasker, and the other *de novo*, with the RepeatModeler2 program.

## I. Introduction

In eukaryotes, most of the genome does not code for proteins or RNA, being made up of non-coding regions (regulatory sequences), introns and repetitive DNA, present in several copies in the genome. Within this last group are the transposable elements (TEs), which are DNA sequences that have the ability to move from one location to another along the genome (Muñoz M. & García JL., 2010). Furthermore, they occupy a large fraction of many eukaryotic genomes (> 80% of the genome of plants such as corn), and have the potential to influence the evolutionary trajectory of their host in three different ways, according to Feschotte C. & Pritham E. (2007): through alterations of genetic function through insertion; by inducing chromosomal rearrangements; or as a source of coding and non-coding material that allows the appearance of novelty (such as new genes and regulatory sequences).

Thus, transposable elements are powerful drivers of genome evolution in many eukaryotes, increasing, over the years, the scientific evidence that transposons contribute to genetic variability, especially in the face of stress conditions during, for example, host-microbe interactions between plants and their associated pathogens, contributing to co-evolution (Seidl MF. & Thomma BP., 2017). Faced with a substantial fraction of plant genomes occupied by TEs, these can participate in various evolutionary processes, such as that of the sex chromosomes of plants, in the suppression of recombination, the diversification of the structure and morphology of the sex chromosomes, sex chromosome degeneration, and dose compensation (Li S. et al., 2019). Thus, the identification and annotation of TEs from dioecious plants would lay the groundwork for further research on the evolution of sex chromosomes.

Within the *Solanaceae* family, studies have been carried out in the analysis of TEs, at the genome level, in *Nicotiana Repandae Section* (Parisod C. et al., 2012); *Calibrachoa (C. parviflora, C. pygmaea* and *C. excellens), Petunia (P. axillaris and P. exserta*) and *Fabiana imbricata* (Kriedt RA. Et al., 2013); *Nicotiana tabacum* (Xu S. et al., 2017); *Solanum lycopersicum* (Jouffroy O. et al., 2016); *Solanum tuberosum* (Jacobs JM. Et al., 1995); and other species (Vicient CM. & Casacuberta JM., 2017). Of all the species mentioned, *Solanum tuberosum* stands out due to being a world-important crop plant that produces high yields of nutritionally valuable foods in the form of tubers. And, at the genomic level, the study of their TEs allows us to better identify and understand the main drivers of the stress response and the evolution of their genome.

Given its importance, the development of clear and efficient TEs annotations has become essential, leading to the use of a variety of software packages that can integrate information from a combination of various TEs detection tools into a composite library, such as TEdenovo. (Grouper, Recon, Piler, LTRharvest) in the REPET package and RepeatModeler (Recon, RepeatScout, TRF) currently dominate the field of TEs identification. Likewise, other tools can search for overrepresented repetitions in unassembled sequence reads, using k-mer counts, machine learning, and low coverage assemblies (Arkhipova IR., 2017).

## II. Hypothesis

The genomes of *Solanum tuberosum subsp. andigena* have a high amount of transposable elements, and there are some differences between these elements in the genomes of the 2 localities used.

## III. Objective

Identify and compare the transposable elements of 2 genomes of *Solanum tuberosum subsp. andigena* from 2 different localities.

## IV. Methodology

### Obtaining the genomes

A search was made in the NCBI database of the complete genomes available from potatoes grown in Peru (GCA_009849725.1 and GCA_009849705.1), which were submitted by McGill University, Canada.

### Homology search

The program RepeatMasker version 4.1.1 (http://www.repeatmasker.org/RepeatMasker/), used in computational genomics, was used to identify, classify and mask repetitive elements, including sequences of low complexity and interleaved repeats. RepeatMasker searches for repetitive sequences by aligning the input genome sequence with a library of known repeats, such as Repbase and Dfam (Tarailo-Graovac, et al. 2009) using hidden Markov profile models. These have been used successfully to discover distant homologues of known proteins in large protein databases, and then this same mechanism was passed on to search for TEs in genomic DNA.

### Analysis of data

The results of the program were taken mainly from the summary provided, of which the graphs were made in the R program with a script programmed by the authors. Subsequently, the other important results were processed using a python script also programmed by the authors to obtain only the classes of the transposons and make a Venn diagram.

## V. Results

It was found that the masked regions occupied 45.22% of the Puno potato genome (*Tables 1, 2*) and 43.32% of the Cusco potato genome (*Tables 3, 4*). Regarding the classes of transposons, in both genomes it was found that there was a higher proportion of retrotransposons, where the highest number were the transposons, Gypsy/DIRS1.

**Table 1.** General information of the data from the Puno potato genome.

|  |  |
| --- | --- |
| **File name** | GCA_009849725.1_ASM984972v1_genomic.fna |
| **Sequences** | 310723 |
| **Total length** | 991109043 bp (991109043 bp excl N/X-runs) |
| **GC level** | 34.84% |
| **Bases masked** | 448192988 bp (45.22%) |

**Table 2.** Summary of the data obtained from the Puno potato genome.

|  | *Number of*<br>*elements* | *Length occupied*<br>*(bp)* | *Percentage of*<br>*sequence (%)* |
| --- | --- | --- | --- |
| *Retroelements* | 421162 | 377526110 | 38.09 |
| *SINEs:* | 17940 | 2337996 | 0.24 |
| *Penelope* | 0 | 0 | 0.00 |
| *LINEs:* | 39121 | 16074439 | 1.62 |
| *CRE/SLACS* | 0 | 0 | 0.00 |
| *L2/CR1/Rex* | 0 | 0 | 0.00 |
| *R1/LOA/Jockey* | 0 | 0 | 0.00 |
| *R2/R4/NeSL* | 0 | 0 | 0.00 |
| *RTE/Bov-B* | 25594 | 5697622 | 0.57 |
| *L1/CIN4* | 18012 | 10914072 | 1.10 |
| *LTR elements:* | 364101 | 359113675 | 36.23 |
| *BEL/Pao* | 0 | 0 | 0.00 |
| *Ty1/Copia* | 52768 | 35238633 | 3.56 |
| *Gypsy/DIRS1* | 300903 | 316642502 | 31.95 |
| *Retroviral* | 0 | 0 | 0.00 |
| *DNA transposons* | 205706 | 47030151 | 4.75 |
| *hobo-Activator* | 46331 | 9960804 | 1.01 |
| *Tc1-IS630-Pogo* | 43321 | 7938377 | 0.80 |
| *En-Spm* | 0 | 0 | 0.00 |
| *MuDR-IS905* | 0 | 0 | 0.00 |
| *PiggyBac* | 0 | 0 | 0.00 |
| *Tourist/Harbinger* | 38315 | 7555986 | 0.76 |
| *Other (Mirage, P-element, Transib)* | 0 | 0 | 0.00 |

**Table 3.** Summary of the data obtained from the Cusco potato genome.

|  |  |
| --- | --- |
| **File name** | GCA_009849705.1_ASM984970v1_genomic.fna |
| **Sequences** | 35961 |
| **Total length** | 841416273 bp (778747073 bp excl N/X-runs) |
| **GC level** | 34.83% |
| **Bases masked** | 364501398 bp (43.32%) |

**Table 4.** Summary of the data obtained from the Cusco potato genome.

|  | *Number of elements* | *Length occupied (bp)* | *Percentage of sequence (%)* |
| --- | --- | --- | --- |
| *Retroelements* | 257659 | 306902963 | 36.47 |
| *SINEs:* | 13556 | 1826372 | 0.22 |
| *Penelope* | 0 | 0 | 0.00 |
| *LINEs:* | 26780 | 13001396 | 1.55 |
| *CRE/SLACS* | 0 | 0 | 0.00 |
| *L2/CR1/Rex* | 0 | 0 | 0.00 |
| *R1/LOA/Jockey* | 0 | 0 | 0.00 |
| *R2/R4/NeSL* | 0 | 0 | 0.00 |
| *RTE/Bov-B* | 18394 | 4240771 | 0.50 |
| *L1/CIN4* | 11501 | 9109403 | 1.08 |
| *LTR elements:* | 217323 | 292075195 | 34.71 |
| *BEL/Pao* | 0 | 0 | 0.00 |
| *Ty1/Copia* | 32044 | 28372509 | 3.37 |
| *Gypsy/DIRS1* | 180237 | 256695476 | 30.51 |
| *Retroviral* | 0 | 0 | 0.00 |
| *DNA transposons* | 140045 | 38651193 | 4.59 |
| *hobo-Activator* | 30987 | 8658993 | 1.03 |
| *Tc1-IS630-Pogo* | 30171 | 5751271 | 0.68 |
| *En-Spm* | 0 | 0 | 0.00 |
| *MuDR-IS905* | 0 | 0 | 0.00 |
| *PiggyBac* | 0 | 0 | 0.00 |
| *Tourist/Harbinger* | 26175 | 6135170 | 0.73 |
| *Other (Mirage, P-element, Transib)* | 0 | 0 | 0.00 |

It was observed that only the transposable elements occupied 42.84% of the complete genome in the Puno potato (*Figure 1A*) and 41.06% in the Cusco potato genome (*Figure 1B*).

**Figure 1.**
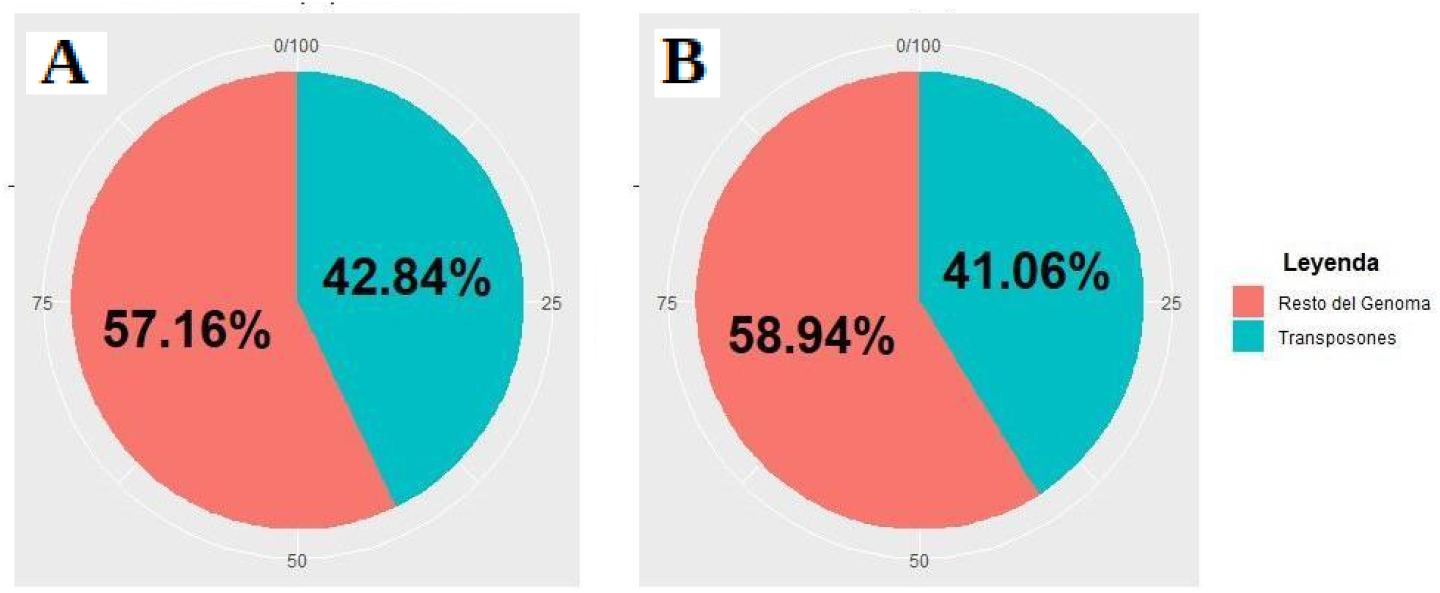
Percentage of transposons in (A) the Puno potato genome, and in (B) the Cusco potato genome.

It was observed that the majority of transposons were class 1, or the so-called retrotransposons and these occupied 88.92% of the complete genome in the Puno potato (*Figure 2A*) and 88.81% in the genome of the Cusco potato (*Figure 2B*).

**Figure 2.**
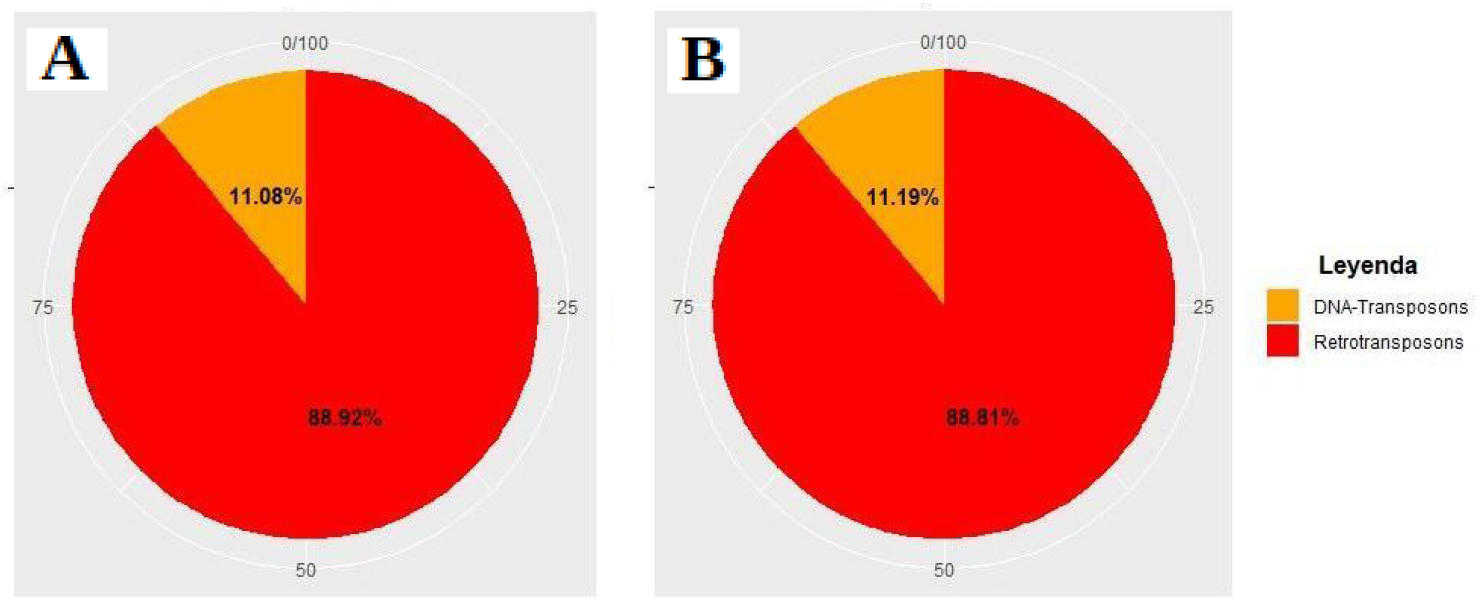
Types of transposons in (A) the Puno potato genome, and in (B) the Cusco potato genome.

It was observed that the majority of transposons were the LTR transposons both for the complete genome in the Puno potato (*Figure 3A*) and for the Cusco potato genome (*Figure 3B*). Followed in quantity by Hobo-Activator transposons and other DNA transposons.

**Figure 3.**
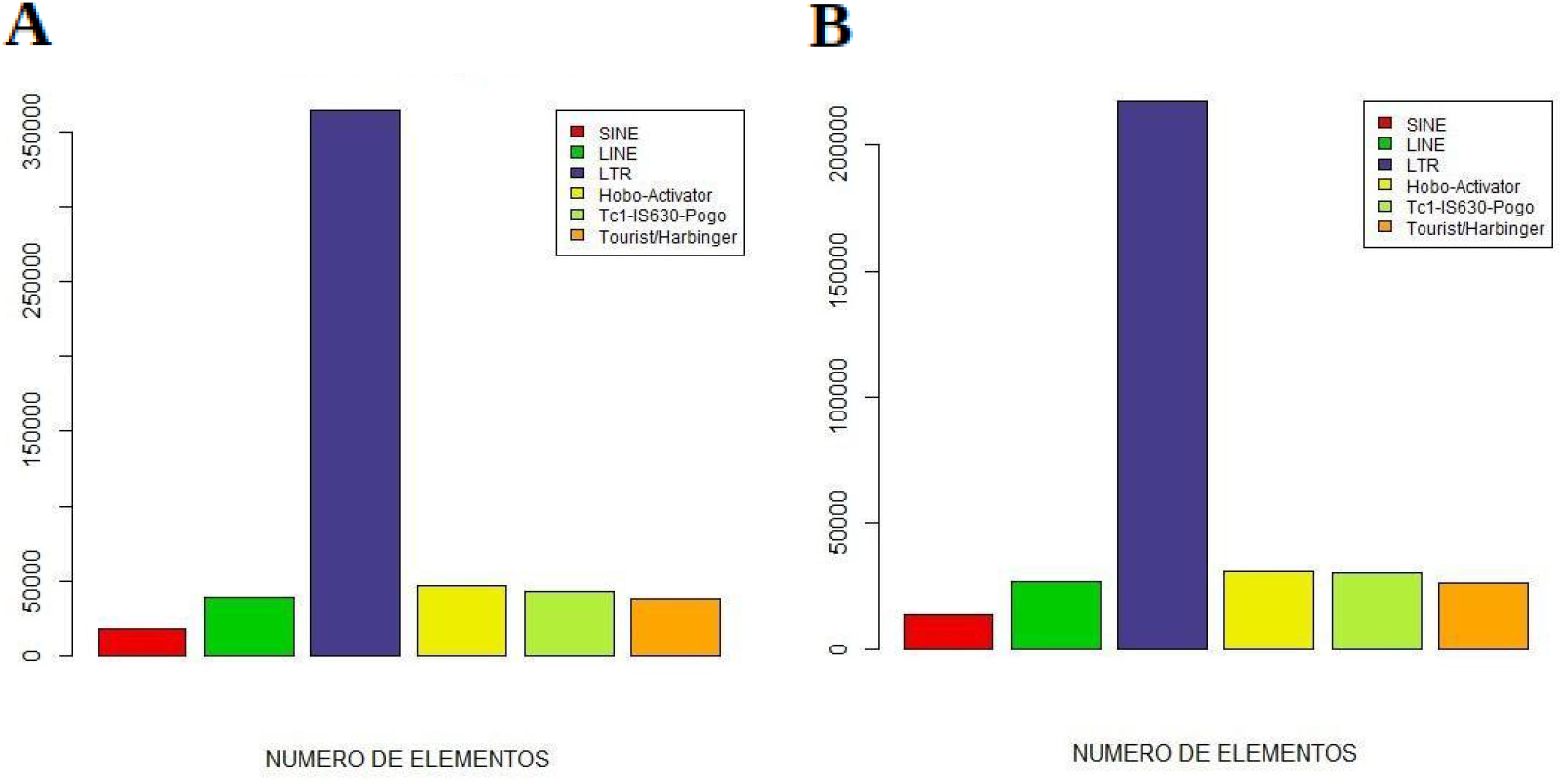
Classes of transposons in (A) the Puno potato genome and in (B) the Cusco potato genome.

It was observed that the LTRs occupied 36.23% of the complete genome in the Puno potato (*Figure 4A*) and 34.35% in the Cusco potato genome (*Figure 4B*).

**Figure 4.**
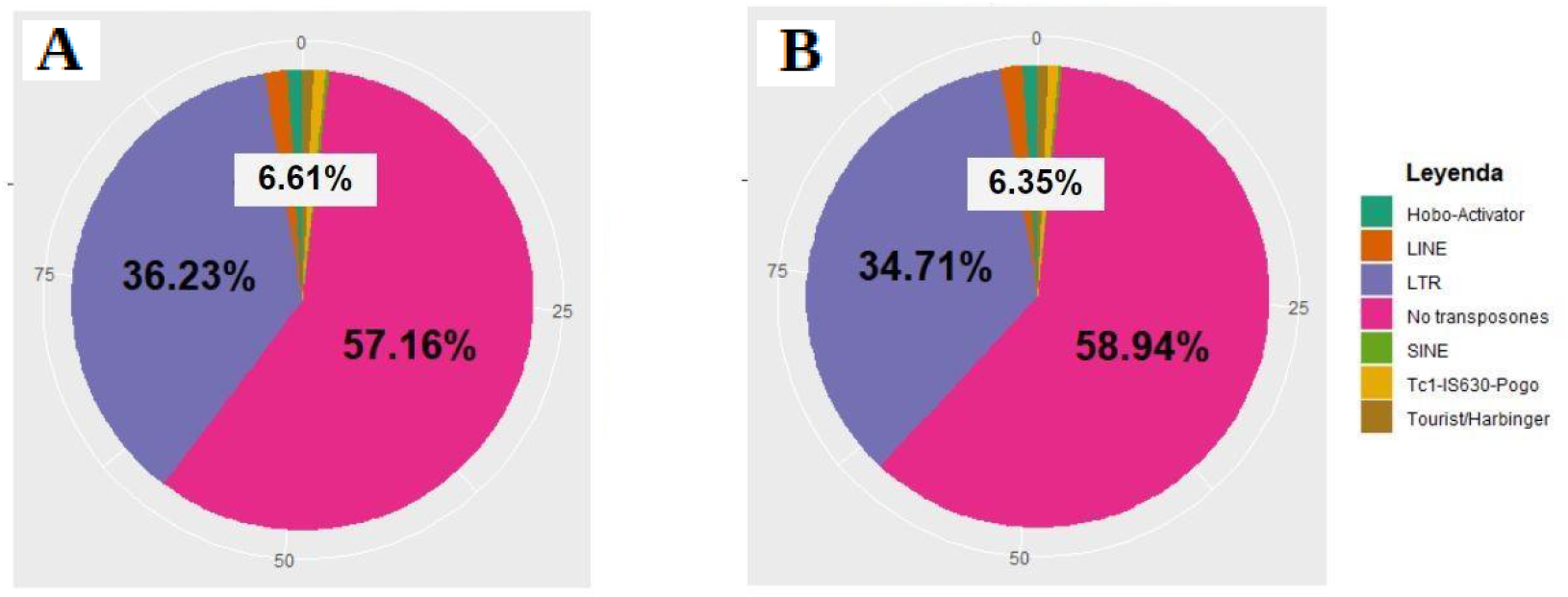
Percentage of transposons in (A) the Puno potato genome and in (B) the Cusco potato genome.

The transposon classes between the two genomes were compared, of which 26 families were found, both for the Puno potato genome and for the Cusco potato genome (*Figure 5*).

**Figure 5.**
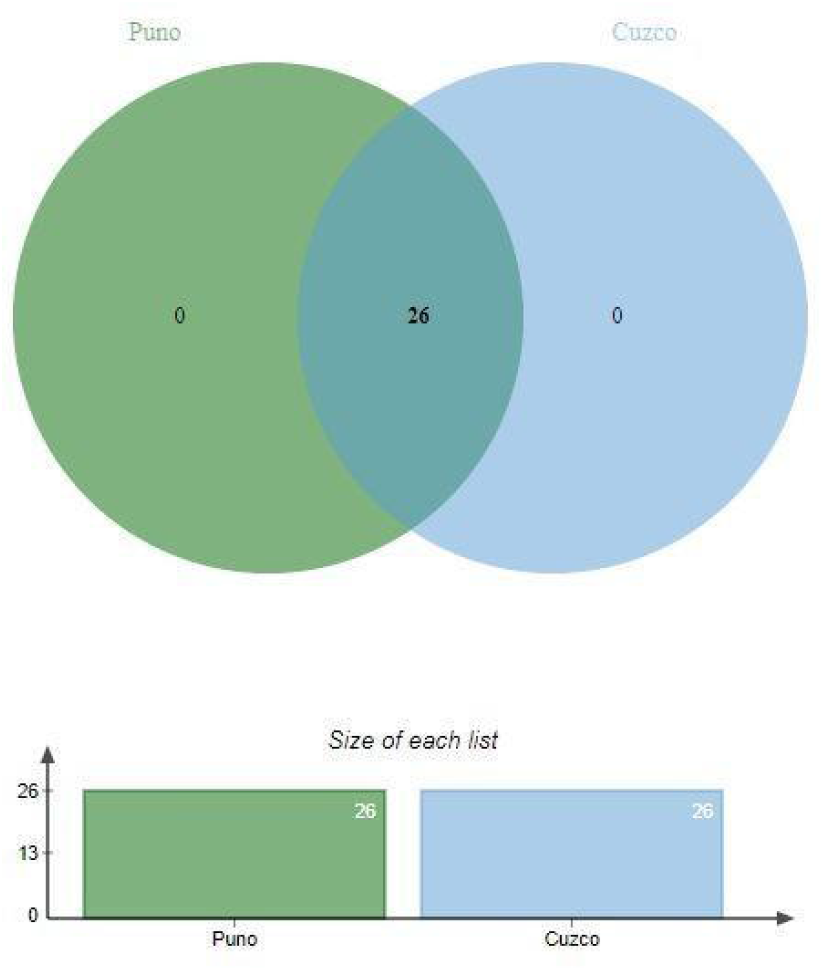
Classes of transposons in Puno and Cusco.

## VI. Discussion

The genus *Solanum* is among the largest plant genera and includes several crops of regional and world importance, among which is the cultivated potato (*Solanum tuberosum*): the third most important food crop after rice and wheat and the main horticultural crop (Devaux et al. 2014).

Currently, the potato in Peru is the main crop in the country in the planted area and represents 25% of the agricultural gross domestic product (GDP). It is the basis of the diet of the Andean area and is produced by 600 thousand small agricultural units, according to what is registered by the Ministry of Agrarian Development and Irrigation (MINAGRI) of Peru. Of all the regions of Peru, Puno is where the most potatoes are planted, harvested and produced, representing 19% and 16% of the total harvested and produced, respectively, in the country during 2017 (IDEXCAM, 2018).

Despite the presence of adverse climate factors (summer, frost, hailstorms in the mountains and excess rain with the presence of pests in the jungle), and the one that is vulnerable to these climatic changes, since 94.56% of its lands Farms are under dry land (MINAGRI, 2008), it maintains a high production of agricultural products (potato, barley grain, quinoa and cañihua). In the same way, at the level of other regions, such as Cusco, they are in the process of growth in relation to the production of potatoes, presenting more than 1,200 types of native potato, of the 3,200 varieties that exist in Peru (Chani A. & Pfuro W., 2015).

Faced with these adverse conditions, transposable elements have been powerful drivers of phenotypic variation that has been selected during domestication and reproduction programs in plant crops (Lisch D. et al., 2013). At the level of the genus *Solanum*, for example, the tomato presents the transpositions of the *Rider* family of long terminal repeat retrotransposons (LTR), which have contributed to several phenotypes of agronomic interest (fruit shape and color), as well as in the regulation of the transcriptional cascade induced by cold stress, and in the face of saline and drought stress by stimulating the accumulation of *Rider* transcripts (Benoit M. et al., 2019). Likewise, the presence of elements similar to *Rider* has been revealed in other economically important crops such as beets and quinoa.

On the other hand, transposable elements exhibit different methylation patterns that are involved in the different adaptation processes of plants (Cantu D. et al., 2010). For example, a retrotransposon-like sequence (*Zm* MI1) showed demethylation patterns under cold stress in corn roots (Steward N., 2000); as well as severe cold stress leads to a decrease in the methylation state and increases the cleavage rate of a specific transposon (Tam3) in *Antirrhinum majus* (Hashida S.N., 2006).

Similarly, the expression of other genes in potato varieties must also be considered, such as, for example, the number of differentially expressed genes (DEGs) in susceptible potato varieties from a study in Huancavelica, where it was suggested that the frost stress response is correlated with positive gene regulation (Martinez D. et al., 2018).

The genomes obtained from *S. tuberosum* in the NCBI database were assembled up to contigs, so they did not have their separation into chromosomes. The genome sizes were 841 Mb and 991 Mb, and they had a GC percentage of 32.20% and 34.80%.

In the results we obtained that more than 40% of the genomes were transposons. This was to be expected, since in plants, TEs can occupy a large proportion of the genome, representing more than half of the total genomic DNA in some cases. For example, the repetitive content comprises approximately 85% and 88% of the wheat and maize genomes, respectively (Schnable et al. 2009; Appels et al. 2018) which may indicate the relevance of these elements for architecture and genome size (Roessler et al. 2018).

The distribution, quantity, and genome coverage of TEs vary greatly, particularly between plants and animals, where LTRs and non-LTRs (LINE and SINE) are the predominant type of TE, respectively (Chalopin et al. 2015). The content of TEs is highly variable in plants and generally shows a positive correlation with genome size (Zavallo D. et al., 2020). For example, up to 85% of the maize genome or 70% of the Norway spruce genome (Nystedt et al. 2013) has been annotated as transposons, including unclassified and other repeating components, while in the more compact genome of *Arabidopsis thaliana* the TEs content is only 21% (Ahmed et al. 2011). In potato, the data presented by Mehra M. et al. (2015) repetitive elements comprised almost 51% of the genome, and when only the most complex elements (i.e. transposons) were considered, the percentage of coverage dropped to almost 34%. A percentage not too far from our larger genomes, where in Puno it was 42.84% and in Cusco it was 41.06%. However, in the study by Zavallo D. et al. (2020) found a TEs content of 18%, which represents half of the genomic coverage according to the data presented by Mehra M. et al. (2015) and ours.

However, of the 18% of the transposons found by Mehra M. et al. (2015), the LTRs comprised around 15% of the potato genome, which corresponds to more than 80% of the total TEs, which was consistent with our results, which was approximately 85% in both genomes. The most abundant superfamily was Gypsy, while the other types of TEs only made up 1% of the genome coverage, also agreeing with our results.

To further understand the discrepancy between TE genome coverage previously reported by Mehra M. et al. (2015), ours and that of Zavallo D. et al. (2020) we must observe the differences between the jobs. For example, in the work of Zavallo D. et al. (2020) although they describe all the types of families of TEs reported by Wicker T. et al. (2007), including LARD, TRIM and MITE, which were absent in the study by Mehra and ours, scored less than half of the sequences, since they applied a clustering method to reduce redundancy and establish a set of filters. selected specifically for each type of TEs, whose objective is to reduce false signals (Zavallo D. et al., 2020). In addition, they used a unique tool that relies on a strategy based on repeatability, leaving aside a wide variety of detection methods that we have combined in this work, which is indicated to improve the detection efficiency of TE (Arensburger P. et al., 2016).

## VII. Conclusions

Based on the results obtained and the discussions carried out, it is presented that, as expected, more retrotransposons are found in the potato genome. Furthermore, almost half of the genomes were transposons.

## Notes

### Competing Interest Statement

The authors have declared no competing interest.

